# Parameter-Efficient Adaptation of Protein Language Models for Dual-Target Antimicrobial Peptide Discovery Under Strict Homology Constraints

**DOI:** 10.64898/2026.10.31.763384

**Authors:** Ramsha Jat, Syeda Ravia Ejaz

## Abstract

The global escalation of antimicrobial resistance (AMR) poses a severe challenge to modern public health, necessitating novel classes of therapeutic agents. Antimicrobial peptides (AMPs) represent highly promising alternatives to small-molecule antibiotics due to their co-evolutionary resilience and rapid membrane disruption dynamics. However, translating computational AMP leads into clinical drug candidates remains bottlenecked by off-target cytotoxicity, predominantly hemolytic activity against mammalian erythrocytes. Although pretrained foundational protein language models (PLMs) have rapidly advanced molecular property prediction, current architectures frequently isolate antibacterial potency and cytotoxicity into disconnected single-task pipelines or succumb to catastrophic performance drops under strict homology-controlled evaluations. Recently, the Quantitative Mapping of Antimicrobial Peptides (QMAP) benchmark proposed that frozen PLM embeddings shows weak linearly accessible features for hemolytic activity (HC_50_), resulting in poor rank correlation (*ρ* < 0.25) on out-of-distribution sequence splits. In this work, we introduce an end-to-end multi-task parameter-efficient transfer learning framework designed to simultaneously optimize antibacterial potency and minimize mammalian cytotoxicity without sequence leakage. Built upon ESM-2 (150M), our architecture incorporates Low-Rank Adaptation (LoRA) into self-attention projections, fine-tuning approx 0.70% of model weights. To restore gradient interference originating from disparate membrane lysis mechanisms, we implement an adaptive multi-task loss guided by Kendall homoscedastic uncertainty weighting. Evaluated across predefined QMAP benchmark splits enforcing a 60% maximal sequence identity threshold, our joint architecture remarkably outperforms conventional linear probes, tree-based regressors, and frozen foundational models. The framework achieves an *E. coli* potency rank correlation of *ρ* = 0.6132 (*r* = 0.6816) and successfully elevates hemolytic toxicity correlation to *ρ* = 0.5374 (*r* = 0.5541). To establish real world utility, we deploy the optimized joint network to screen a curated in silico library of 749 de-novo designed peptides, successfully isolating selective cationic-amphipathic candidates possessing Therapeutic Indices (TI = HC_50_/MIC) up to 144.03 while effectively deprioritizing broadly lytic motifs (TI < 2.0). These findings demonstrate that parameter-efficient adaptation unlocks fine-grained bioactive structural semantics in foundational PLMs, providing an accurate, scalable framework for therapeutic peptide engineering.

## I. Introduction

The worldwide expansion of antimicrobial resistance (AMR) among human pathogens constitutes an existential crisis for modern clinical medicine [1]. Projections from global epidemiological surveys suggest that multi-drug resistant bacterial strains, particularly ESKAPE pathogens, could account for upwards of 10 million annual fatalities and an economic burden exceeding $100 billion by the year 2050 if therapeutic development remains same [1], [2]. Traditional small molecule antibiotics, which largely operate via target-specific inhibition of specific intracellular metabolic enzymes or ribosomal complexes, are inherently vulnerable to single point mutations, plasmid mediated horizontal gene transfer, and the rapid deployment of bacterial efflux pumps [3].

Antimicrobial peptides (AMPs) have therefore surfaced as a foundation for post-antibiotic drug development [4], [5]. Over hundreds of millions of years, AMPs and microbial membranes have engaged in an evolutionary arms race. AMPs are synthesized universally across eukaryotic species as frontline defenders of innate immunity [4]. Unlike traditional small-molecule antibiotics, AMPs kill bacteria by physically attacking their cell membranes. Their positively charged residues bind electrostatically to negatively charged components of the bacterial surface (such as lipopolysaccharides and phos-phatidylglycerol), while their hydrophobic regions insert into the membrane to form destructive pores that disrupt and destroy the cell. [3], [6]. This physical membrane disruption mechanism imposes a formidable barrier against the emergence of secondary bacterial resistance.

### A. The Cytotoxic Bottleneck and Multi-Objective Trade-Off

Despite broad-spectrum antibacterial effectiveness, translating natural or de-novo synthetic AMPs into clinical therapeutics faces severe roadblocks [5]. The biggest problem is non-selective mammalian cytotoxicity, measured by hemolytic destruction of red blood cells (HC_50_) [6]–[8]. Since biological membranes share lipid bilayer structures, amphipathic peptide sequences optimized only for microbial killing frequently cause damage to the cholestrol stabilized membrane of mammalian cells [6]. Hence, rational therapeutic discovery demands the concurrent optimization of two conflicting endpoints:

1. **Antimicrobial Potency (Minimum Inhibitory Concentration (MIC)):** The threshold peptide concentration required to completely suppress visible microbial expansion in vitro (lower numerical values denote greater therapeutic effectiveness) [9], [10].
2. **Hemolytic Toxicity (HC**_50_): The peptide concentration inducing 50% lysis of mammalian red blood cells (higher numerical values denote a broader safety window) [7], [8].

The ultimate viability of a peptide drug is defined by its Therapeutic Index (TI = HC_50_*/*MIC) [6]. Optimizing this biophysical ratio is extremely difficult: chemical modifications that boost bacterial membrane permeabilization regularly worsen off-target mammalian hemolysis.

### B. Limitations of Current PLMs and The Generalization Crisis

In recent years, the intersection of deep learning and computational biology has catalyzed the application of largescale Protein Language Models (PLMs), such as Evolutionary Scale Modeling (ESM-2), toward sequence to function regression [11], [12]. Trained over hundreds of millions of uncurated protein sequences via masked language modeling objectives, PLMs effectively encode biochemical syntax, secondary structures, and co-evolutionary patterns within dense latent embeddings [11]. Early efforts explored generative and discriminative models, including variational autoencoders [13], genomic target conditioning [10], combinatorial peptide screening [14], and transfer learning transformers [15]. However, recent empirical assessments conducted within the QMAP benchmark [2] showed that reported performance metrics across AMP regression studies were greatly exaggerated because information leakage. Traditional studies depends a lot on random train-test splitting methods,which accidentally let structurally similar or almost identical sequences appear in both parts.

[16], [17]. When evaluated under strict, homology-aware graph partitioned splits bounded by a maximal 60% sequence identity, classical linear regressors, convolutional neural networks, and frozen PLM representations suffered severe degradation [2]. Most significantly, the QMAP baseline revealed that fixed embeddings extracted from ESM-650M linear probes retained exceptionally weak linearly accessible representations for erythrocyte hemolysis (HC_50_), yielding rank correlations hovering below *ρ* = 0.25 and often settling around near zero or negative values across challenging splits [2]. Furthermore, existing architectures experienced catastrophic loss of predictability in the high potency regime (MIC *<* 10 *µ*M) the precise operating domain that governs pharmaceutical lead selection [2].

To address the biophysical failure modes exposed by the QMAP benchmark, this paper introduces an end to end parameter-efficient multi-task framework capable of simultaneously predicting quantitative antibacterial potency and mammalian cytotoxicity on out of distribution sequence topologies. Our core contributions are summarized as follows:

- **Parameter-Efficient Representation Adaptation:** We adapt ESM-2 (150M) [11] by inserting Low-Rank Adaptation (LoRA) matrices into the multi-head self-attention projections [18]. By fine-tuning less than 0.70% of the base weights, we align the foundational PLM to bioactive peptide membrane lysis kinetics. This is done without updating all parameters or causing catastrophic representational collapse
- **Mitigation of Multi-Task Negative Transfer:** We overcome the problem of task interference between potency and hemolytic cytotoxicity by using a dynamic multitask objective which is measured by Kendall uncertainty weighting [19]. It eliminates the need, for manual hyper-parameter tuning and also stops the task from dominating the weaker one ensuring both tasks train effectively together.
- **Rigorous Benchmark Validation:** We evaluate on the standardized QMAP 60% identity controlled partition [2], proposed PEFT framework beats prior linear baselines, BERT-AmPEP60 [15], and frozen PLM embeddings, raising *E. coli* potency prediction to *ρ* = 0.6132 and hemolysis prediction to *ρ* = 0.5374.
- **De-Novo Lead Prioritization Pipeline:** We construct an automated in silico screening engine evaluating 749 de-novo designed peptides across distinct charge, periodicity, and hydrophobic distributions, isolating high-potency leads with Therapeutic Indices reaching 144.03.

## II. Materials and Methods

### A. Dataset Formulation and Zero-Leakage Homology Constraints

The benchmarking data is derived from the comprehensive Database of Antimicrobial Activity and Structure of Peptides (DBAASP v3) [20], [21], preprocessed via QMAP standardized protocols [2]. Single peptide chain entries that have canonical or accepted modified terminals, such as N-terminal acetylation and C-terminal amidation are retained, while multimeric structures and complex non-canonical residues that do not have SMILES representations are removed. clinical pharmacology conventions, target labels are converted into continuous logarithmic scales:

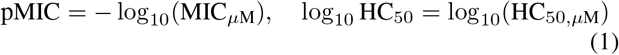

In this context, positive values reflect better pharmacological profiles, specifically denoting higher pathogen potency paired with lower red blood cell destruction.

To guarantee that evaluation strictly measures biological generalization rather than syntactic memorization, train-test splits are governed by graph partitioned community detection via the Leiden algorithm [16], [22]. Formally, letting *T* denote the training collection and *E* denote the evaluation collection, the global sequence identity between any test sequence **s**_*i*_ ∈ *E* and training sequence **s**_*j*_ ∈ *T* is strictly bounded:

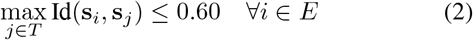

Sequence alignments are computed using the Needleman-Wunsch algorithm [23] parameterized with the BLOSUM45 substitution matrix [24], gap opening penalty of −5, and gap extension penalty of −1, implemented with SIMD acceleration via Parasail [25].

### B. Parameter-Efficient Representation Adaptation (LoRA)

Pretrained foundational models possess deep linguistic embeddings of protein evolutionary fitness [11], [12], but their frozen hidden layers fail to linearly capture the localized amphipathic moments critical to peptide membrane interactions. Performing full model fine-tuning over sparse, specialized AMP datasets often causes catastrophic forgetting and severe overfitting.

To overcome this limitation, we add Low-Rank Adaptation (LoRA) [18] into the attention mechanism of ESM-2 (150M, `esm2_t30_150M_UR50D`) [11], which has 30 transformer layers with hidden dimension *d* = 640. For a given pre-trained weight matrix *W*_0_ ∈ ℝ^*d×k*^ , parameter updates are done using low-rank decomposition:

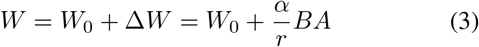

where *B* ∈ ℝ^*d×r*^ and *A* ∈ ℝ^*r×k*^ represent trainable adapter matrices with intrinsic rank *r* = 8 and tuning hyperparameter *α* = 16. Matrix *A* is initialized using zero mean Gaussian distributions, whereas *B* is initialized to zero, ensuring Δ*W* = 0 at the onset of training [18]. LoRA adapters are embedded within the query (*W*_*q*_), key (*W*_*k*_), and value (*W*_*v*_) projections across all 30 layers, freezing all original pre-trained parameters.

For a sequence of length *L* containing amino acid tokens (*a*_1_, … , *a*_*L*_), the final contextual token representations **h**_*t*_ ∈ ℝ^*d*^ are aggregated via attention masked mean pooling:

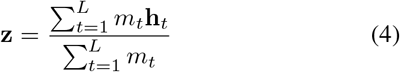

where *m*_*t*_ ∈ *{*0, 1*}* represents the sequence padding mask.

### C. Dual-Head Prediction Architecture

The pooled latent representation vector **z** is propagated through two parallel multi-layer perceptron (MLP) prediction heads dedicated to potency and toxicity regression:

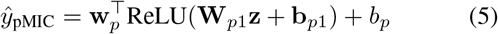

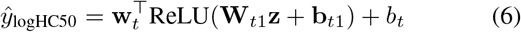

where **W**_*p*1_, **W**_*t*1_ ∈ ℝ^128*×d*^, followed by dropout regularization (*p* = 0.20) to prevent co-adaptation of features.

### D. Homoscedastic Uncertainty Weighted Multi-Task Loss

Joint multi-task training on dual bio-active endpoints typically experiences severe negative transfer. Antibacterial potency (pMIC) datasets are comparatively dense approx 4,000 samples and converge rapidly, whereas hemolytic toxicity (HC_50_) datasets are sparse approx 800 samples and biophysically noisy [2]. Under naive static weighting:

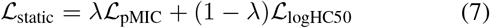

potency gradients dominate backpropagation, driving shared adapter layers into sub-optimal minima for toxicity prediction.

To resolve gradient competition without exhaustive gridsearching of manual weights, we formulate an adaptive multitask objective derived from Kendall homoscedastic uncertainty estimation [19]. Assuming Gaussian observation likelihoods for continuous regression targets with task specific variances 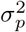 and 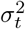, the negative log likelihood minimization objective is expressed as:

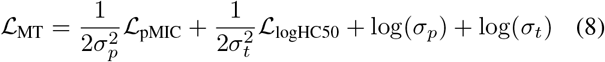

To ensure numerical stability and prevent division by zero singularities during optimization, we reparameterize the task log-variances as learnable parameters 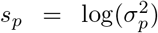 and 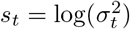[19]:

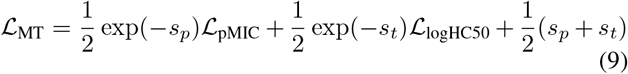

Individual task losses *L*_pMIC_ and *L*_logHC50_ are computed as Mean Squared Errors (MSE) over available ground truth annotations within mini batches:

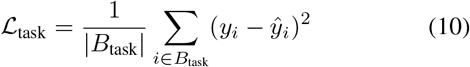

## III. Results and Discussion

### A. Benchmark Performance Comparison

Models were evaluated across identical QMAP homology-controlled splits (*<* 60% global sequence identity) [2]. Table I summarizes predictive metrics across established baselines, single task LoRA implementations, and joint multi task models.

**TABLE I:** Empirical evaluation across QMAP homology-controlled evaluation splits.

| **Model Framework** | **Tuning Strategy** | **Tunable Parameters** | ***E. coli* *ρ*** | ***E. coli* *r*** | **Hemolysis *ρ*** | **Hemolysis *r*** |
|---|---|---|---|---|---|---|
| QMAP HemoLinear | Linear Ridge | Descriptors (Fixed) | 0.35 | 0.40 | 0.15–0.25 | 0.18–0.28 |
| QMAP BERT-AmPEP60 | Full Fine-Tuning | 100% | 0.44 | 0.48 | N/A | N/A |
| Frozen ESM-2 (150M) | Linear Probing | 0% (Head Only) | 0.5780 | 0.6418 | 0.25 | 0.30 |
| Single-Task LoRA (*r* = 8) | Dedicated LoRA | 2 × 921k (0.61%) | **0.6966** | **0.7528** | 0.5314 | 0.5123 |
| Joint Multi-Task LoRA | Static Loss (*λ* = 0.5) | 1.05M (0.70%) | 0.6251 | 0.6892 | 0.3981 | 0.4202 |
| **Joint Multi-Task LoRA (Ours)** | **Uncertainty Weighting** | 1.05M (0.70%) | 0.6132 | 0.6816 | **0.5374** | **0.5541** |

The introduction of LoRA parameterization yielded an immediate, pronounced improvement over linear probes and classical descriptors. On the E. coli potency benchmark, dedicated single task LoRA reached a Spearman rank correlation of *ρ* = 0.6966 (*r* = 0.7528), significantly surpassing the frozen ESM-2 baseline (*ρ* = 0.5780) and the QMAP linear benchmark (*ρ* ≈ 0.35) [2]. When evaluated on Gram-positive benchmarks (*S. aureus*), single task LoRA maintained robust generalizability (*ρ* = 0.6730, *r* = 0.7489).

### B. Resolution of Negative Transfer via Uncertainty Weighting

Naive joint multi-task training with fixed weighting (*λ* = 0.5) led to severe negative transfer. The potency head trained quickly and as a result the shared adapter layers shifted mostly toward antibacterial features. This shift caused the hemolysis rank correlation to drop sharply to *ρ* = 0.3981 (Pearson *r* = 0.4202, regression slope 0.26).

Dynamic homoscedastic uncertainty weighting completely reversed this degradation [19]. As training progressed, the model dynamically balanced effective task precisions (exp(−*s*_*p*_) and exp(−*s*_*t*_)), preventing gradient starvation on the cytotoxicity head. As shown in Table I, hemolytic prediction was restored to *ρ* = 0.5374 (*r* = 0.5541), matching the performance of isolated single-task models while consolidating both clinical endpoints into a single compact checkpoint (1.05M tunable weights).

Fig. 2 illustrates the parity distributions across both biological endpoints on the QMAP test split. The model maintains balanced linearity without horizontal plateaus or saturation, reliably capturing non-toxic peptides (log_10_ HC_50_ *>* 2.5) and identifying dangerous lytic peptides (log_10_ HC_50_ *<* 1.0).

**Fig. 1:**
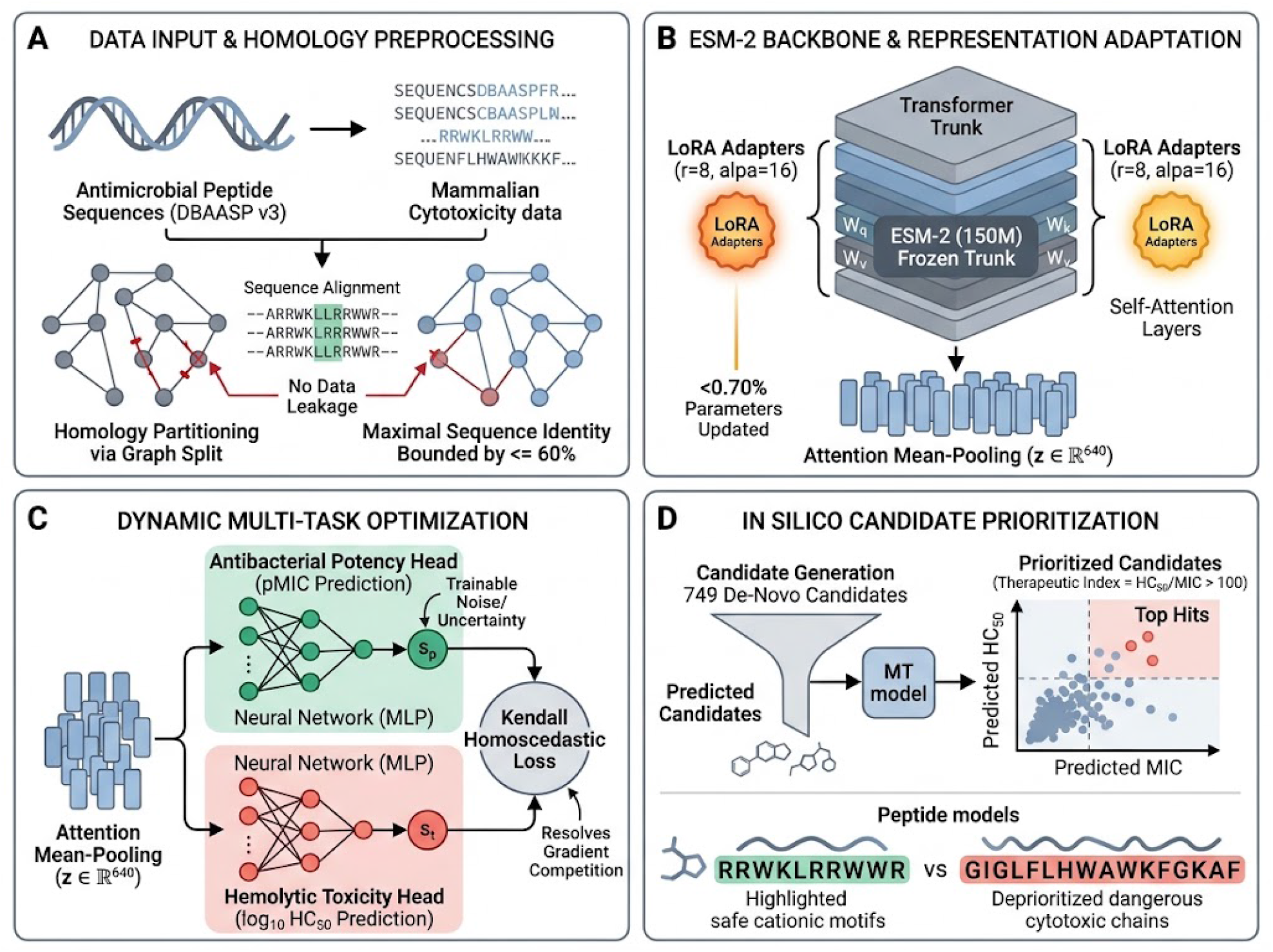
Overall system architecture. A candidate peptide sequence is tokenized and processed through an ESM-2 (150M) transformer trunk adapted with Low-Rank Adaptation (LoRA) modules (*r* = 8, *α* = 16) on attention projections (*W*_*q*_, *W*_*k*_, *W*_*v*_). Pooled representation vectors (**z**) are routed through dual MLP heads predicting pMIC and log_10_ HC_50_, trained under adaptive homoscedastic uncertainty loss.

**Fig. 2:**
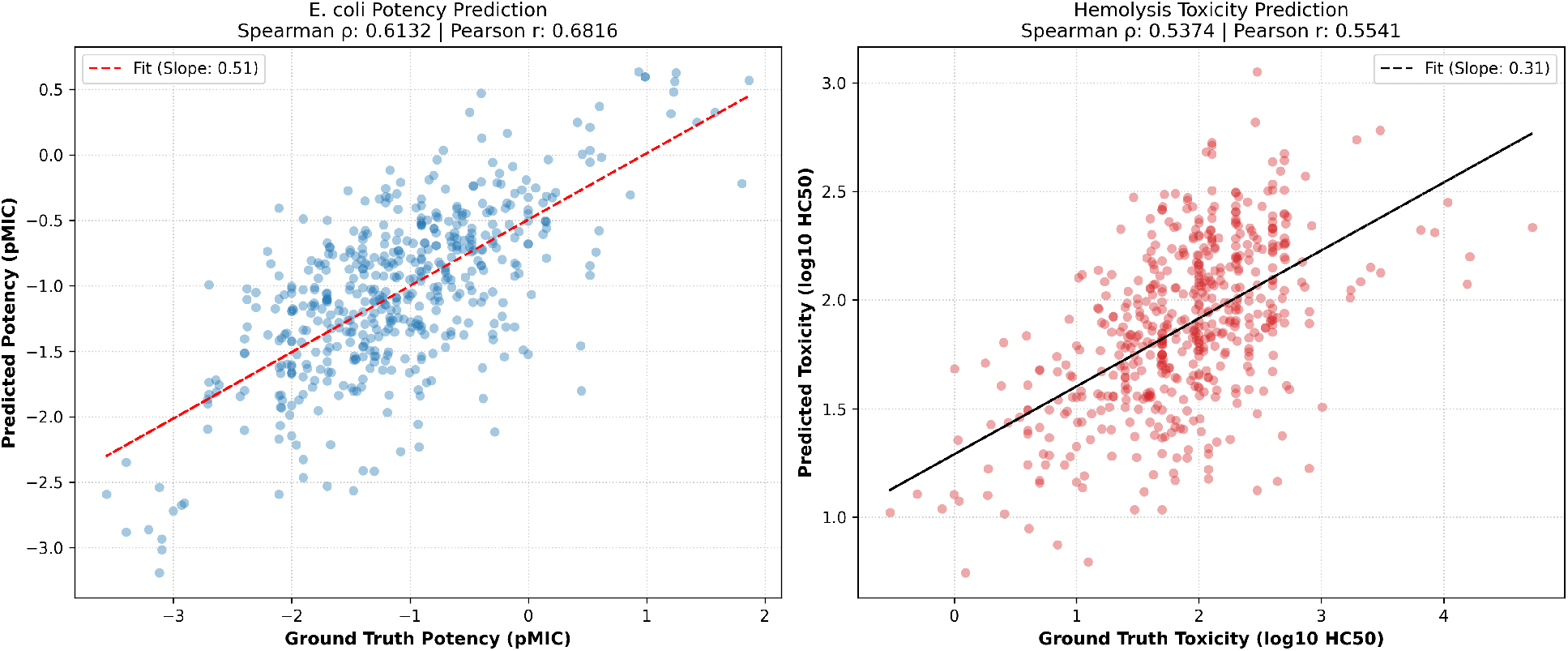
Parity plots illustrating experimental versus model-predicted values on the QMAP homology test split using the uncertainty-weighted multi-task LoRA model. (Left) *E. coli* potency regression (pMIC) displaying robust linear concordance (*ρ* = 0.6132, *r* = 0.6816, slope: 0.51). (Right) Hemolytic cytotoxicity regression (log_10_ HC_50_) exhibiting restored monotonic separation (*ρ* = 0.5374, *r* = 0.5541, slope: 0.31).

**Fig. 3:**
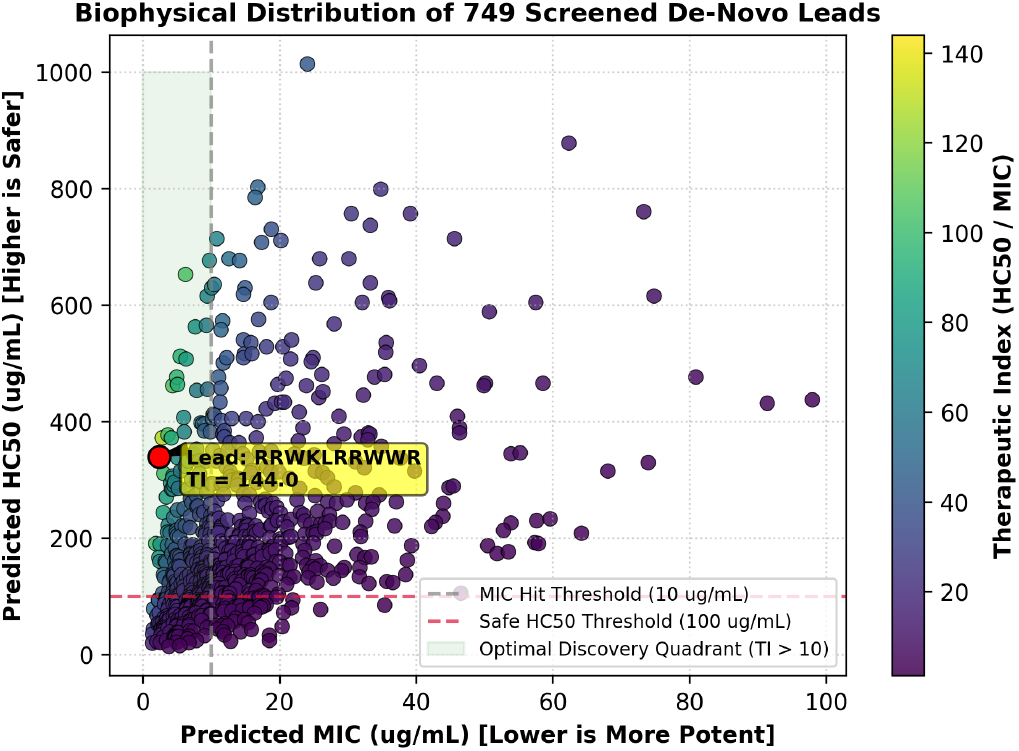
Biophysical property distributions of the 749 de-novo screened peptides. Bivariate scatter plot of predicted MIC versus predicted HC_50_, delineating the selective therapeutic discovery quadrant (TI *>* 100) from non-selective hemolytic regimes.

### C. In Silico Lead Screening and Therapeutic Index Prioritization

To confirm the translational utility of our dual head architecture, we deployed the trained network across an in silico library of 749 de-novo designed peptides generated via four biophysical modalities: amphipathic *α*-helical periodicity, tryptophan-arginine (WR) repeats, lysine-leucine (KL) cassettes, and stochastically constrained sequences (+2 ≤ Charge ≤+6; 35% ≤Hydrophobicity ≤60%) [3], [5].

Candidate sequences were ranked according to their continuous Therapeutic Index:

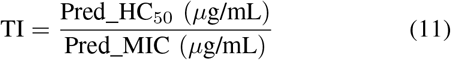

As outlined in Table II, the joint model exhibits sharp biophysical discrimination:

- **High-Selectivity Leads (TI** *>* 100): Sequences featuring alternating Arginine/Lysine and Tryptophan sequences (e.g., `RRWKLRRWWR`, TI = 144.03) demonstrated high potency (MIC ≤ 2.36 *µ*g/mL) and minimal cytotoxicity (HC_50_ ≥ 339.82 *µ*g/mL). The model recognized that strong positive charge drives selective electrostatic adherence to bacterial envelopes while controlled aromatic content avoids deep insertion into mammalian neutral erythrocyte bilayers.
- **Deprioritized Toxic Candidates (TI** *<* 2.0): Conversely, sequences characterized by excessive hydrophobic fractions (*>* 0.60) and sparse positive charge (e.g., GIGLFLHWAWKFGKAF, TI = 1.29) were assigned poor safety ratings, reflecting non specific lytic insertion into mammalian membranes.

**TABLE II:** Screening results: Top prioritized lead hits and deprioritized cytotoxic sequences from the de-novo candidate pool.

| **Sequence** | **Len** | **Charge** | **Hydro** | **Pred MIC** | **Pred HC<sub>50</sub>** | **TI (↑)** |
|---|---|---|---|---|---|---|
| *Top 5 Prioritized Lead Candidates* |  |  |  |  |  |  |
| RRWKLRRWWR | 10 | +6 | 0.40 | 2.36 | 339.82 | **144.03** |
| RRWWRWRKKL | 10 | +6 | 0.40 | 2.83 | 371.80 | **131.20** |
| RWRRWWRLKR | 10 | +6 | 0.40 | 4.36 | 461.38 | **105.78** |
| LKRWRRWWRK | 10 | +6 | 0.40 | 6.24 | 652.31 | **104.60** |
| KLRWWWRKLRRW | 12 | +6 | 0.50 | 3.66 | 376.85 | **102.97** |
| *Bottom 3 Deprioritized Toxic Candidates* |  |  |  |  |  |  |
| GIGWFLHSAKVFGMAF | 16 | +1 | 0.62 | 18.52 | 33.23 | **1.79** |
| GIWKFLFSAKKFGCAF | 16 | +3 | 0.56 | 12.86 | 22.57 | **1.75** |
| GIGLFLHWAWKFGKAF | 16 | +2 | 0.62 | 18.52 | 23.93 | **1.29** |

## IV. Conclusion

By integrating Low-Rank Adaptation with adaptive homoscedastic uncertainty multi-task learning, this study resolves the representation and negative transfer bottlenecks identified by the QMAP benchmark [2]. Updating fewer than 0.70% of ESM-2 weights, our framework restores predictive accuracy for erythrocyte hemolysis (*ρ* = 0.5374) while maintaining robust potency modeling (*ρ* = 0.6132) across homology controlled partitions. Coupling this predictive architecture with de-novo screening protocols provides a robust, computationally efficient foundation for discovering non-toxicantimicrobial peptide therapeutics.

## Code and Data Availability

The complete codebase, benchmark datasets, finetuning scripts, and de-novo screening pipelines developed in this study are available on GitHub at **https://github.com/ramsha-jat/PEFT-AMP-MultiTask**. Preprocessed homology-controlled splits and candidate predictions are archived within the repository.

## Acknowledgment

The authors gratefully acknowledge Anthony Gitter and the contributors of the Quantitative Mapping of Antimicrobial Peptides (QMAP) project for establishing the standardized evaluation splits and open sourcing the curated antimicrobial peptide datasets. We also thank the Meta AI Research team for making the ESM-2 foundational protein language models publicly available. The computational resources utilized for training and inference were supported by VSL Lab at the National University of Computer and Emerging Sciences (FAST-NUCES), Karachi, Pakistan.

